# Biologically plausible gated recurrent neural networks for working memory and learning-to-learn

**DOI:** 10.1101/2023.07.06.547911

**Authors:** Alexandra R. van den Berg, Pieter R. Roelfsema, Sander M. Bohte

## Abstract

The acquisition of knowledge does not occur in isolation; rather, learning experiences in the same or similar domains amalgamate. This process through which learning can accelerate over time is referred to as learning-to-learn or meta-learning. While meta-learning can be implemented in recurrent neural networks, these networks tend to be trained with architectures that are not easily interpretable or mappable to the brain and with learning rules that are biologically implausible. Specifically, these rules employ backpropagation-through-time for learning, which relies on information that is unavailable at synapses that are undergoing plasticity in the brain. While memory models that exclusively use local information for their weight updates have been developed, they have limited capacity to integrate information over long timespans and therefore cannot easily learn-to-learn. Here, we propose a novel gated recurrent network named RECOLLECT, which can flexibly retain or forget information by means of a single memory gate and biologically plausible trial-and-error-learning that requires only local information. We demonstrate that RECOLLECT successfully learns to represent task-relevant information over increasingly long memory delays in a pro-/anti-saccade task, and that it learns to flush its memory at the end of a trial. Moreover, we show that RECOLLECT can learn-to-learn an effective policy on a reversal bandit task. Finally, we show that the solutions acquired by RECOLLECT resemble how animals learn similar tasks.

## Introduction

A hallmark of human intelligence is the capacity to accumulate knowledge over different learning experiences. This capacity not only allows the acceleration of learning in the same domain, but can also facilitate learning in similar domains, a phenomenon referred to as learning-to-learn (Harlow, 1949; Thrun & Pratt, 1998). Standard neural network models lack this ability and quickly forget previously acquired knowledge when they are trained on a new task, a phenomenon called catastrophic forgetting (French, 1999). This is particularly problematic in the case of reversal learning (Izquierdo et al., 2017), where two stimuli are initially associated with a certain reward probability (e.g. stimulus A with a 75% chance of reward and stimulus B with a 25% chance of reward), but then reverse their stimulus-reward associations (stimulus A is rewarded with 25% probability and stimulus B with 75% probability). In this case, the network has to fully change its weight structure to compensate for these changes. To overcome this limitation, meta-learning models have been developed that over the course of several similar tasks acquire a set of weights that facilitate generalisation to novel tasks which bear similarities to those they had been trained on.

Meta-learning can be achieved using a multitude of approaches (Wang, 2021; Sutton, 2022; Huisman, van Rijn, & Plaat, 2021). The most promising approach from a biological perspective uses recurrent neural networks trained with reinforcement learning (Wang et al., 2018; Duan et al., 2016). These networks are trained on a distribution of tasks, where they are provided with information about previous actions, rewards and observations. Subsequently, they are tested on tasks from the distribution that had not yet been seen, while the weights of the network are fixed. If meta-learning has taken place, it should be able to perform this task even in the absence of new learning. In this framework, the algorithm used to train the network is able to give rise to a new algorithm nested in network activations, rather than its weights, that can efficiently learn new tasks in a way that bears similarity to how the prefrontal cortex together with the basal ganglia and thalamus might come to learn-to-learn (Wang et al., 2018). In other words, this property would enable networks to identify and store in memory (as an activity pattern) which task it is performing and to switch the memory pattern when the task changes. Hence, task switching can happen within one trial by changing the activity pattern in memory as opposed to going through the elaborate process of retraining the entire network connectivity.

While their behaviour is more similar to biological systems, a limitation of these meta-learning models is that the learning rules and architectures used for this purpose are biologically implausible. In particular, these recurrent networks tend to be trained using backpropagation-through-time (BPTT), which relies on information that is not available at the level of the synapse for its updates (i. e. it is non-local in time; Lillicrap & Santoro, 2019). An example of an algorithm that solves this temporal credit assignment problem is e-prop (Bellec et al., 2021). E-prop approximates BPTT by using synaptic traces. These are local signals that are stored within synapses that monitor and record their contribution to the current activity in memory units. An earlier example of a memory network that used a local update rule involving synaptic traces is AuGMEnT (Rombouts et al., 2015), which additionally solves the spatial credit assignment problem by using attentional feedback signals from selected actions (i.e. tags) to select which synapses are responsible for an action and should be updated. However, this model remembers sensory information by accumulating it over time and does not have the gating mechanisms required to flexibly update memories by choosing which information to retain or forget. To compensate for this, the network parameters in AuGMEnT are reset frequently to flush the entire memory. As a result, it can only integrate information over relatively short timeframes, which hinders its ability to learn-to-learn.

Whether training is done with a local learning rule or not, the specific gated architecture (i.e. long-short term memory, or LSTM) often used for meta-learning can also be questioned from a biological perspective. The LSTM relies on three multiplicative gates that control the information contained in memory cells (Hochreiter & Schmidhuber, 1997). However, the full LSTM architecture can be more complex than what is actually required to solve various tasks (Ravanelli et al., 2018; Dey & Salem, 2017). As a result, it becomes difficult to interpret this model and map it to the brain. Simplifications to the LSTM have been proposed, such as the gated recurrent unit (GRU) with only two gates (Cho et al., 2014), and more recently, the light-gated recurrent unit (Light-GRU) with a single gate (Ravanelli et al., 2018). These models are not only more interpretable, but have also been shown to yield good or even superior performance on some tasks compared to the standard LSTM architecture (Cho et al., 2014; Ravanelli et al., 2018).

In this study, we propose RECOLLECT: a novel light-gated recurrent unit that is exclusively trained using information that is both local in space and time, making it biologically plausible. RECOLLECT adapts the synaptic tags and traces from AuGMEnT (Rombouts et al., 2015) to implement a learning rule for the Light-GRU architecture that closely approximates BPTT. We show that RECOLLECT can flexibly use its working memory to perform a pro-/anti-saccade task and learns-to-learn on a reversal bandit task. Finally, we illustrate similarities between how RECOLLECT and animals acquire these tasks.

## Results

### Architecture

#### Feedforward processing

The aim of this study is to develop an architecture that can learn to forget in a manner that is plausible for biological systems. Specifically, the architecture itself should be mappable to the brain and it should be trained using a learning rule where all the information necessary for the synaptic change is available locally, at the synapse.

The novel model is called “REinforCement learning of wOrking memory with bioLogically pLausible rECurrent uniTs” - RECOLLECT (see Figure 1). RECOLLECT draws inspiration from two models: the light-gated recurrent unit (Light-GRU; Ravanelli et al., 2018), and AuGMEnT (Rombouts et al., 2015).

**Figure 1.**
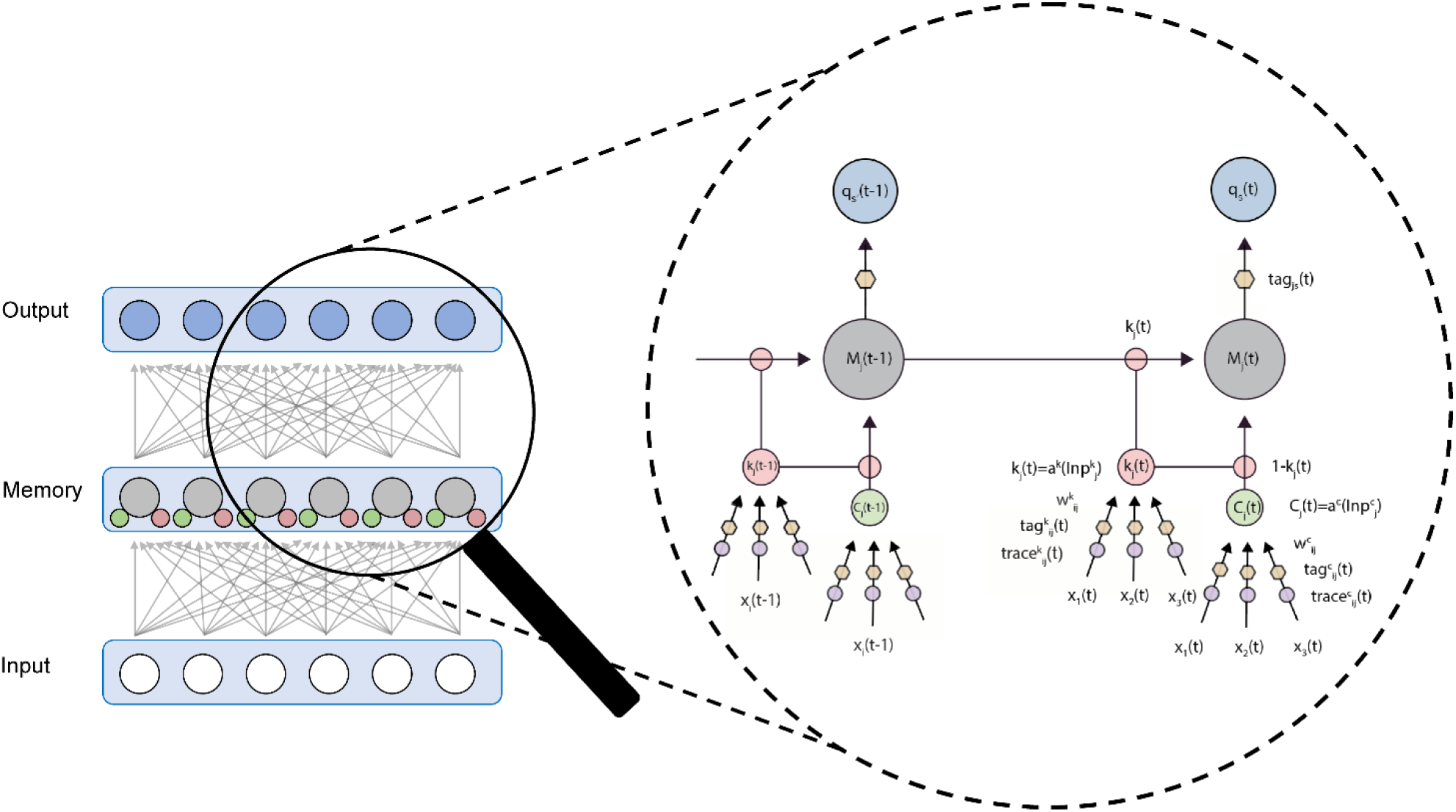
RECOLLECT Architecture. Gating units (k_j_) balance the extent to which past memories are retained versus how much novel information from candidate input cells (C_j_) can enter memory. The resulting memory units (M_j_) synapse onto output unitsthat estimate Q-values of actions. Synaptic tags and traces store information that is necessary for the synaptic updates.

The Light-GRU (Ravanelli et al., 2018) is a recurrent network that processes information from the current sensory environment, as well as from the same network at the previous timestep; its memory. Information flow is regulated by a learnable ‘gate’ that determines the degree to which these two sources are considered. Consider the case of a sequential task where a new number needs to be added to a cumulative sum of previous numbers until it is told to stop. For the first timestep, the memory content will be 0. If the number presented is 2, the network will output 2. For the second timestep, the memory content is 2, which is added onto the new number 5 to output 10. However, if on the next timestep the new sensory information consists of the number 7, but also a ‘stop’ signal, then the memory content is reset to 0 and the network will only output 7. In the case of a more complex task, the gate may have a more sophisticated way to balance old and new information. It can be speculated that such a network in the brain can correspond to a loop involving the cortex, thalamus, basal ganglia or even cerebellum (see Discussion).

RECOLLECT consists of an input layer, a memory layer with GRUs and an output layer. As in the Light-GRU (Ravanelli et al., 2018), the memory layer contains three types of units: candidate input cells (*C_j_*), gating units (*k_j_*) and memory cells (*M_j_*), which can be considered as being located in the same cortical column. Current sensory information (*x_i_*(*t*)) is processed by the candidate input cells and available to update the activity of the memory cell.

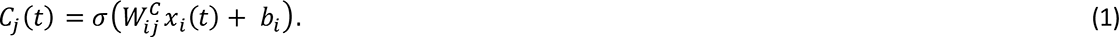

Here, *C_j_*represents the activity of the candidate input units, 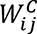 denotes the synaptic weights, *b_i_* the bias and *σ*(·) is the sigmoidal activation function used to constrain outputs of units between 0 and 1.

The gating units determine the degree to which past and new information together form the updated memory. This is done by controlling the degree of forgetting of the state of the memory unit, and the relative contribution of the candidate input cells that deliver new information. The activity of the gating units, *k_j_*, depends on the input through weights 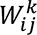:

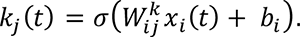

The gating units determine the updating of the activity of memory units *M_j_* as follows:

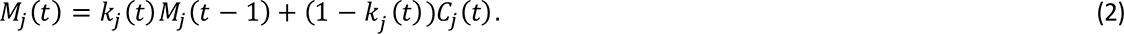

If gating units are active, the candidate input cells do not have much influence on the memory unit and the previous memory *M*_*j*_ (*t* − 1) is retained. Vice versa, if the gating units are weakly activated, the memory units will make a large step in the direction of the activity level *C*_*j*_ of the candidate input cells. We therefore refer to this gate as a memory gate. The process by which RECOLLECT uses memory gates to balance memorisation and forgetting is depicted in Figure 2A.

**Figure 2.**
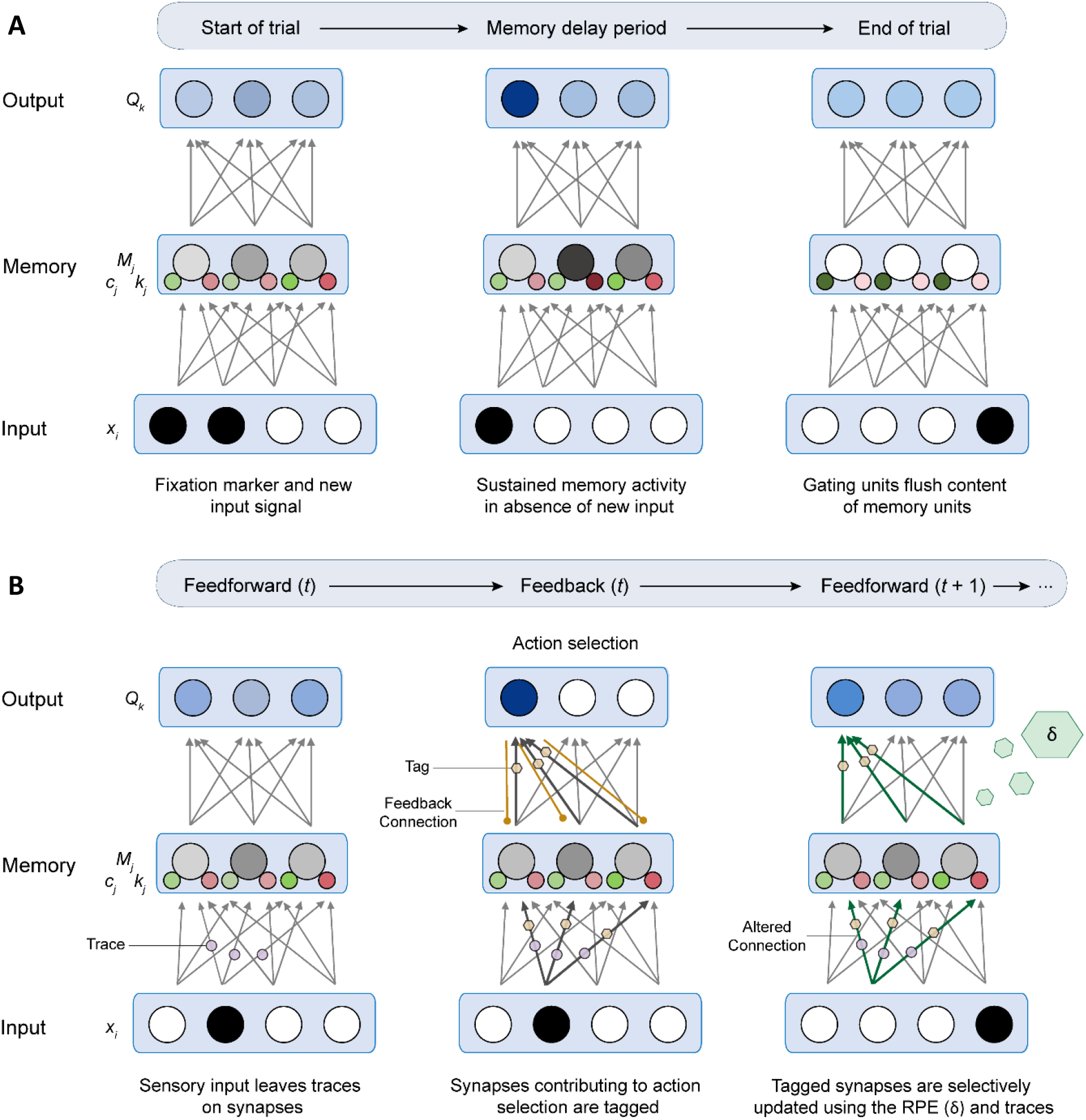
A) RECOLLECT flexibly selects when input should be remembered or forgotten. Memory units increase their activity when new sensory information is acquired. This activity can be sustained over a memory delay if the gating activity is high. When an input signalling theend of a trial is shown, gating units decreasetheir activity and thereby allow the memory units to forget the information they have accumulated over the trial. B) Formation of synaptic tags and traces in RECOLLECT. Activation of input units during feedforward processing can create synaptic traces on connections to gating and candidate input units. Upon action selection, relevant synapses contributing to theselected actions are tagged by means of feedback connections. A reward prediction error in the form of a global neuromodulator is released when the expected reward based on the Q-value of the selected action is different from theactual reward that is received. These tagged synapses areeither potentiated or depressed depending on the sign of the reward prediction error. If the reward is better or worse than expected, the tagged connections are potentiated and depressed, respectively.

Once the memory units have been updated, the activity of the memory units is propagated to the output units.

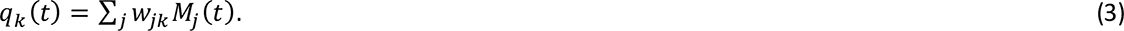

The output units estimate the Q-value *q_k_*, the expected (discounted) reward of each action *k* that can be taken by the network. Once these values have been computed, an epsilon-greedy strategy selected the winning action *s*, where the action with the highest Q-value is chosen with probability 1-ε, and a random action is selected with probability ε.

#### Learning rule

RECOLLECT defines a learning rule for the Light-GRU that is based on synaptic tags and traces and relies exclusively on information local to the synapse. This rule is equivalent to BPTT when the model does not use recurrent weights (as in the model described in the previous section) and a close approximation thereof if these weights would be included (see Method). In this section, we explain the equations outlining this learning rule.

Once RECOLLECT selects an action *s*, it may receive a reward. If this reward differs from the expected reward based on the Q-value of the chosen action, this discrepancy can be captured in the form of a reward prediction error (RPE, *δ*):

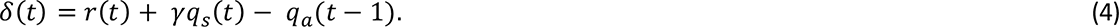

This computation follows the SARSA temporal difference learning rule, which compares the predicted outcome of the previous action q*_a_*(*t*-1) to the sum of the observed reward *r(t)* and the discounted Q-value of the winning unit *q_s_*(*t*). A negative RPE indicates that the outcome was worse than anticipated, whereas a positive RPE signals that a higher reward was received than was estimated at the previous time step. The RPE is presented to the network in the form of a global neuromodulator, hence it is a signal that is accessible for all synapses in the network.

When synapses are exposed to the RPE, plasticity can occur. As in AuGMEnT (Rombouts et al., 2015), plasticity is regulated using tags and traces. Tags are eligibility traces (Sutton & Barto, 2018; Houk, Adams & Barto, 1995) that are formed on all synapses that contributed towards action selection and register how much a synapse contributed to the selected action. To do this, they require feedback from the selected action to the memory layer. After their formation, the tags interact with the global neuromodulator that provides information about the RPE. Consequently, only those synapses that were tagged will become plastic.

Aside from the tags, synaptic traces are maintained on connections from input units to the candidate input units and gating units. These traces store presynaptic activity in the aforementioned synapses over time. Specifically, if a specific input unit *i* contributed to the activation of a memory unit *j*, then the trace_ij_ keeps track of how much of this input is still visible in the memory activity, even if this input was given several time steps in the past.

Using a combination of these tags and traces, all the information that is required for network updates is locally available (see Figure 2B for a schematic illustration of the learning rule). The following equations will define the updates for the tags, traces and weights for each of the units in RECOLLECT.

For the output units, the tags are formed in the presence of both presynaptic activity (*M*_*j*_) and postsynaptic activity after an action is selected (*z*_*k*_). The *Tag*_*jk*_ only increases if the output unit *k* is selected, i.e. if *k*=*s*, in which case the presynaptic activity *M_j_* of memory unit *j* is added onto the tag:

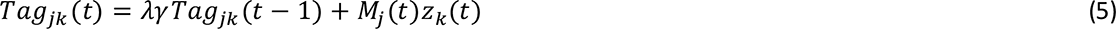

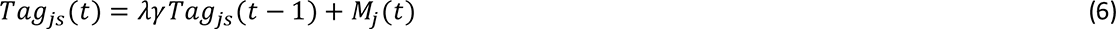

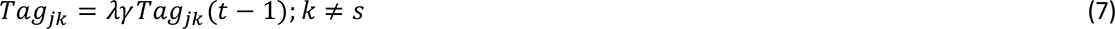

Once a tag is formed, it decays according to two hyper-parameters: the tag decay rate (*λ*) and the reward discount factor (*γ*). As a result, synapses contributing to previous actions can still be affected by network updates in subsequent timesteps, but to a smaller extent as time progresses. This learning scheme corresponds to the temporal difference (λ) algorithm (Sutton, 1988).

The weight update Δ*w*_*jk*_ depends on the tag, the RPE and the learning rate (*β*):

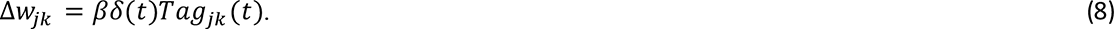

Aside from the tags, the update of the synapses 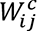 from the candidate input cells to the memory units also depend on the degree to which these input cells contributed to 0-value *q_s_*of the selected action *s*. Their influence is indirect, through the memory unit *j.* Plasticity therefore depends on (i) how much unit *j* contributed to the Q-value of the selected action and (ii) on the contribution of this synapse on the memory unit *j*’s activity level on the current and previous time steps. RECOLLECT (as in AugMEnT; Rombouts et al., 2015) uses a ‘trace’ to keep track of the synapse’s influence on the activity level of memory unit *j* and a ‘tag’ to estimate the influence of the synapse on the Q-value of *s*. We start with describing the update of the trace, which is initialized at a value of 0:

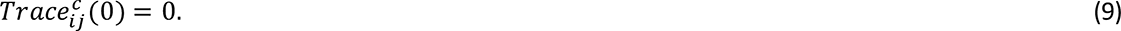

The influence of the synapse 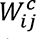 on the activity of memory unit *j* depends on the slope of the activation function 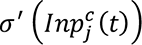 of the input unit at time *t*, the activity of the input unit *x_i_*, and negatively on the activity of the memory gate, which together define the second term in the equation below. The first term represents the influence of the synapse on the activity of memory unit *j* on previous time steps 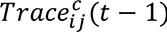:

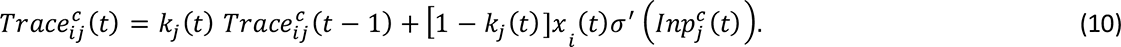

Note that this contribution quickly declines if *k*_*j*_ (*t*) is small, i.e. if the memory gate is biased towards updating the activity of the memory unit.

It is now also possible to estimate the influence of this synapse on the *q_s_*, based on information that is available locally in the synapse because it was stored in the trace, as follows:

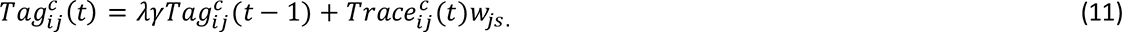

The second term includes *w_js_*, which equals the feedback that arrives at the memory unit *j* through the feedback connection from the winning output unit *s*. This attentional feedback signal is proportional to the contribution of unit *j* to the Q-value of the selected action. Furthermore, the first term permits TD(*λ*) in case *λ* is larger than 0, as was described above for the weights between the memory units and the output layer.

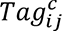 interacts with the learning rate *β* and the globally released neuromodulator that signals *δ* to form the weight update, as was also described above:

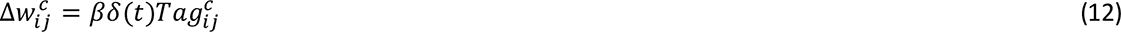

We will now consider the plasticity of the connections of the gating units, which are updated equivalently, using tags and traces. The trace is initialized at time 0:

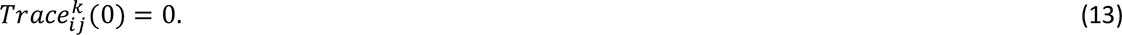

The contribution of the synapse 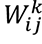 to the activity of the memory unit *j* depends on the slope of the activation function 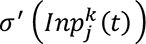, the activity of the input unit *x_i_* and the difference between the activity of the memory unit at the previous time step *M_j_*(*t*-1) and the new input to the memory unit *C_j_*, which is intuitive because the setting of the memory gate is irrelevant if the activity of the candidate input is equal to the activity of the memory unit on the previous time step. The first term in the equation below represents the influence of the synapse on the activity of the memory unit on previous time steps 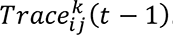.

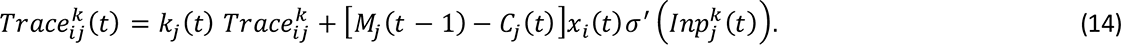

The equations for the tag and the weight update are equivalent for those of the connections to the candidate input units:

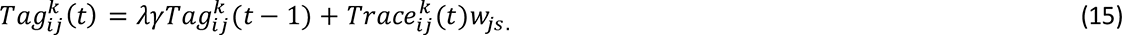

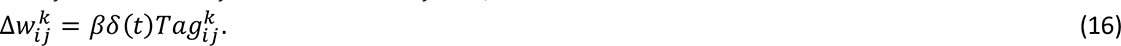

Compared to output units, the main addition is that synaptic traces are used for updating the tag. These traces are first initialised at 0, after which they start accumulating presynaptic activity for every subsequent timestep.

#### Biological plausibility

In addition to using only local information in its learning rule, RECOLLECT has various other properties that were inspired by neurobiology. For instance, the output units in RECOLLECT encode for Q-value of actions. Neurons coding for action values have been observed in several regions, including the midbrain (Morris et al., 2006), basal ganglia (Hikosaka, 2014; Ito & Doya, 2009) and frontal cortex (Rushworth et al., 2011; Cai & Padoa-Schioppa, 2014; Padoa-Schioppa & Assad, 2006).

Moreover, to shape plasticity RECOLLECT makes use of a global neuromodulatory signal that conveys the reward prediction error. Such prediction errors are believed to be generated by midbrain dopamine neurons and support decision-making and learning (Schultz, 2016). Other than the reward prediction error, there is also the sensory prediction error. This has been described as a signal comparing the actual versus the intended sensory feedback and is believed to drive learning as well (Keller & Mrsic-Flogel, 2018). Equation 14 incorporates a comparison between the old memory unit and the current candidate input unit [*M*_*j*_ (*t* − 1) − *C*_*j*_ (*t*)]. This term relates to the sensory prediction error (see e.g. Keller & Mrsic-Flogel 2018).

The update rule also integrates biological features. Namely, tags (or eligibility traces) are used to demarcate synapses that contribute to the winning unit. Experimental findings have been made to support the existence of these type of traces (Gerstner et al., 2018; Yamaguchi et al., 2022). Furthermore, since only the tagged synapses are updated, learning follows a type of Hebbian plasticity (Magee & Grienberger, 2020) where both presynaptic and postsynaptic activity, in combination with the RPE, shape network updates.

In conclusion, RECOLLECT is a biologically inspired model that is both equipped with a gated memory that allows for selective forgetting and integration of information over longer timespans, as well as with a learning rule that closely approximates BPTT, but exclusively using information that is locally available at the level of the synapse (see proof in Method).

### RECOLLECT selectively gates relevant information into working memory

The goal of the current paper was to develop a memory model that can learn to forget using a local learning rule. To investigate how RECOLLECT gates information into its working memory and how it sustains these memory representations over time, the model was trained on the pro-/anti-saccade task from Gottlieb and Goldberg (1999). This task was previously used to train AuGMEnT (Rombouts et al., 2015): a biologically plausible model of memory, but which cannot forget. As such, this task can be used to illustrate differences in how these models learn.

The task (see Figure 3A) started with an empty visual display, after which either a blue or green fixation marker appeared. If fixation was not acquired within 10 timesteps, the trial was terminated without reward. Otherwise, the model received a fixation reward of 0.2 and was presented with a cue on either the left or the right side of the fixation marker for a single timestep. Once the cue disappeared, a memory delay commenced for 5 timesteps. If fixation was broken at any time before the end of this delay, the trial ended and no more reward was given. If the model kept fixating, the marker was removed and the model had to decide whether to make a saccade to the left or to the right. Only if the model chose correctly within 8 timesteps, it received a reward of 1.5. Importantly, the correct saccade direction depended both on the colour of the fixation marker and on the location of the cue. The blue fixation colour marked pro-saccadic trials where the saccade had to be made in the direction where the cue had been presented, whereas the green fixation colour marked anti-saccadic trials on which the saccade had to be made in the opposite direction.

**Figure 3.**
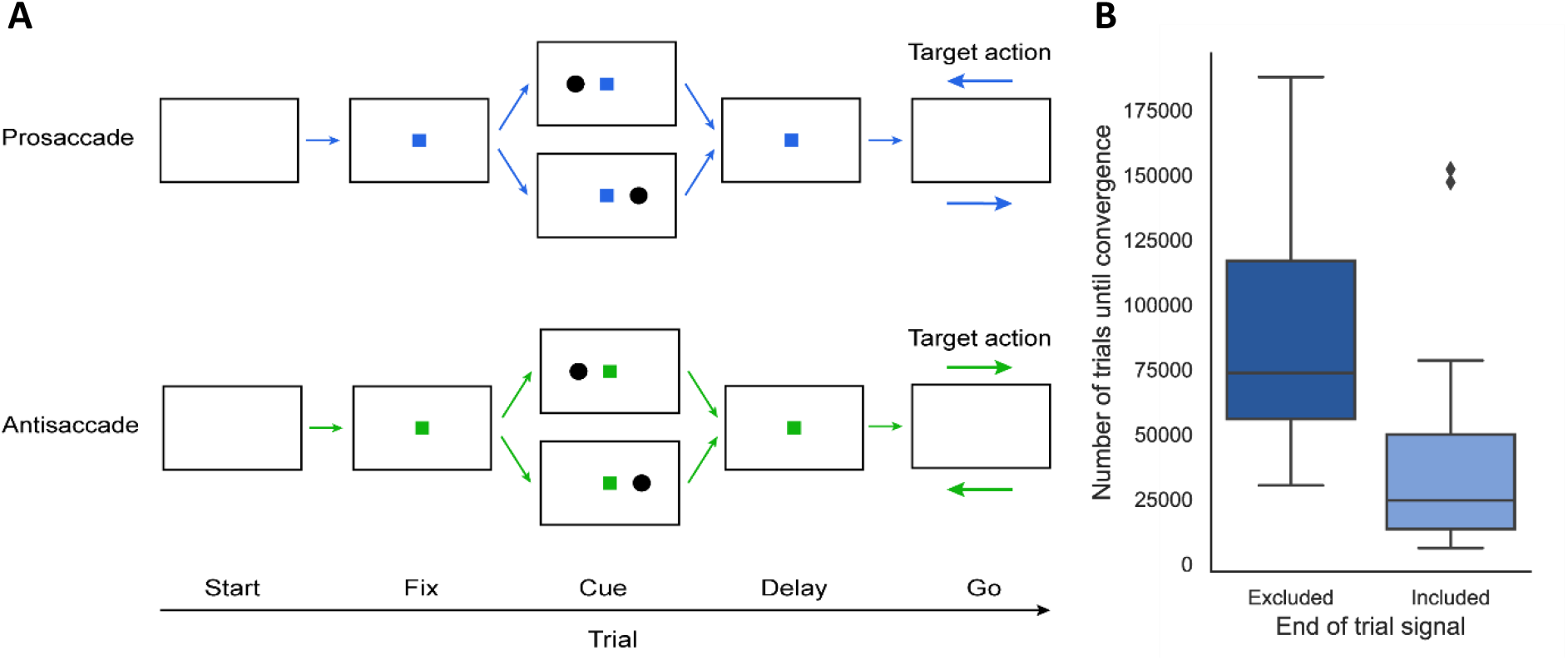
A) Structure of the pro-/anti-saccade task. The fixation colour indicates whether a pro-saccade (blue) or anti-saccade (green) to a cue on the left or right side of the fixation mark has to be performed after a memory delay. The input layer in RECOLLECT receives fixation colour and cue location information. The output layer encodes the three actions that can be taken: fixation, saccade to the left or a saccade to the right. B) Boxplot of the number of trials before convergence when an end of episode signal is included in the input (left) or when it is not (right).

Correct performance on the task thus not only depended on learning the value of both the fixation colour and cue location and performing a non-linear computation thereupon, but also required remembering this information for the entire duration of the delay period, as well as forgetting it upon the end of the trial to prevent interference with subsequent trials.

In order to train RECOLLECT on the task, one-hot encoded input was provided about the two possible fixation marker colours and the left or right cue. The network was trained for a maximum of 1.000.000 trials or until convergence. Convergence was established if 1) the model had reached criterion performance (85% correct trials) on the last 100 trials of the four trial types (i.e. pro-saccade left, pro-saccade right, anti-saccade left and anti-saccade right), and 2) when it could perfectly complete all four trial types with its weights fixed and exploration disabled (i.e. no more learning could occur).

Out of 20 random initialisations (with 4 input units, 7 gating, candidate input and memory units each, and 3 output units) of RECOLLECT that were trained on the pro/anti-saccade task, all networks reached the convergence criterion, indicating that RECOLLECT indeed successfully utilised its working memory. However, more training was required before convergence than in our previous AuGMEnT model when using a comparable network size^1^. Specifically, the median number of training trials required was 73614 for RECOLLECT, but only 4100 for AuGMEnT. An important difference between RECOLLECT and AuGMEnT is the extent to which memory is maintained. AuGMEnT remembers which stimuli were presented on the task by default and this memory is artificially flushed at the end of every trial. In reality, organisms have to learn the structure of repetitive events and also that information that is relevant during a trial may be forgotten at the end of it, because these memories can interfere with subsequent trials. In our experiments, RECOLLECT had to learn which information it needed to retain and when it should be forgotten. The comparison with AuGMEnT reveals that such a more versatile gating mechanism requires additional training time.

We hypothesized that learning could be accelerated if we would add explicit cue indicating the termination of a trial because the network might learn to use this signal as a right moment to flush its memory. Indeed, the inclusion of this end-of-trial signal reduced the median number of trials before convergence from 73614 to 24657 trials (Wilcoxon signed-ranks test for 20 randomly initialised networks with and without reset signal indicated a large effect size, *z* = -2.99, *p* = 0.003, *r* = 0.67) (Figure 3B).

To investigate how RECOLLECT solves the pro-/anti-saccade task, we examined the tuning of the units. To create more insight into how these units bridged the memory delay and intertrial interval, a curriculum was used to extend these timesteps until 5 and 3 timesteps, respectively (see Materials & Methods). Figure 4 shows the activity of units from an example network for the four trial types. While this network converged already at the 11331^th^ trial, training was continued until the 15000^th^ trial to highlight differences in selectivity between trial types.

**Figure 4.**
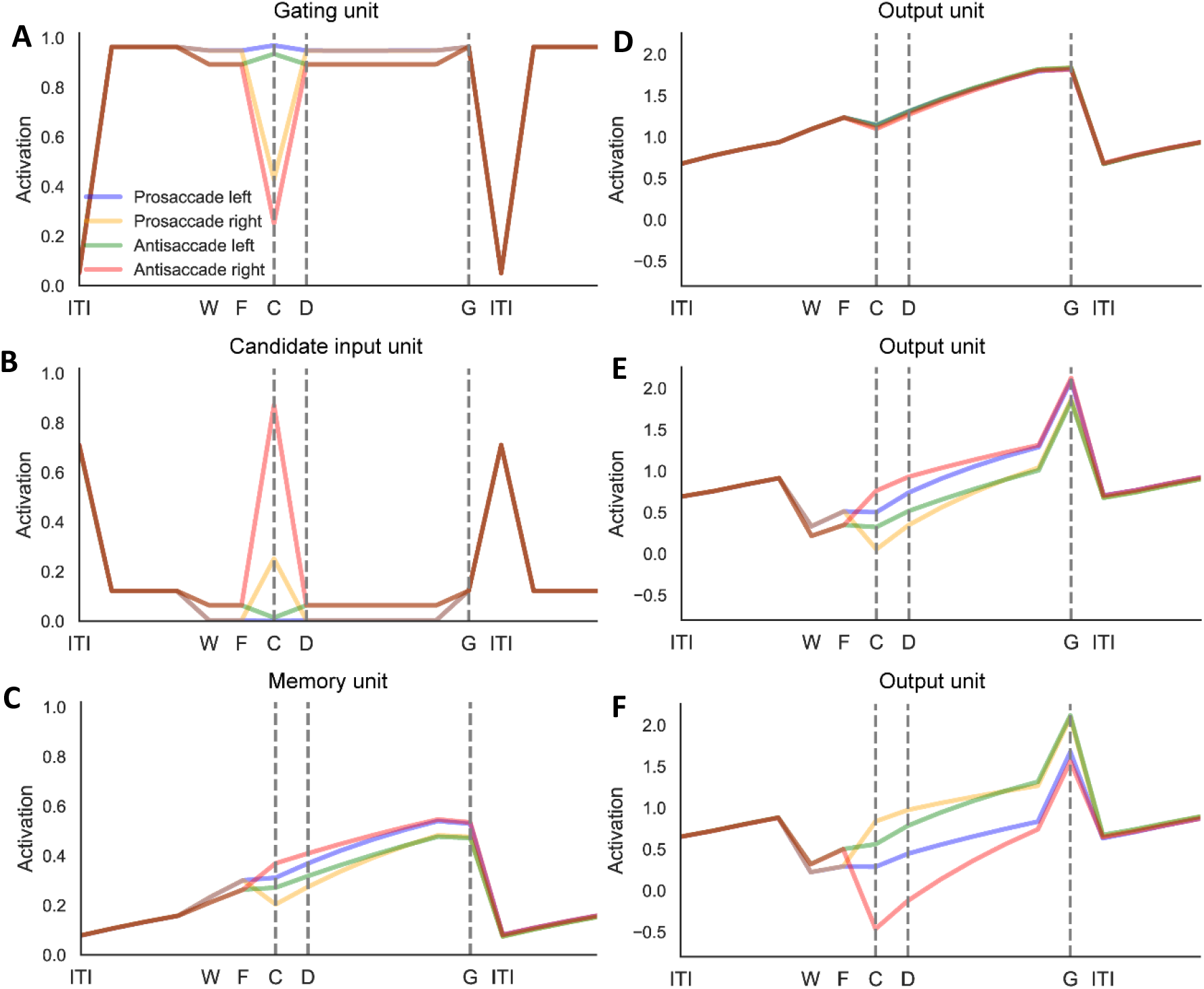
The activity of example units in a network trained on the pro-/anti-saccade task. A) Gating unit that was sensitive for cues on the left side, with strong activity on pro-saccadic trials. B) Candidate input unit that responded to right cues on anti-saccadic trials. C) Memory unit with a weak bias for leftward saccades. D-F) The output units approximating the q-value of fixation (D), a saccade to the left (E) and to the right (F). Labels: ITI = intertrial interval, W = waiting period until fixation is acquired, F = time of fixation, C = cue presentation, D = memory delay period, G = go.

Units developed selectivity for the type of saccade (pro- or anti-saccadic), the location of the visual cue (left or right), to combine these two types of information to select the appropriate saccade. For instance, the gating unit illustrated in Figure 4A responded to left cues and was slightly more active on pro-saccadic trials. In some cases, units were selective for a single trial type, such as the candidate input unit in Figure 4B, which was strongly active for anti-saccadic trials with a cue on the right. Several memory units developed selectivity for the required saccade direction, coding for the appropriate eye movement during the memory delay both on pro-saccade and anti-saccadic trials. For instance, the memory unit in Figure 4C displayed a slight selectivity for leftward eye movements. As required by the task, the output unit with the highest Q-value was the one coding for the required action. (Note that the Q-values for the non-coding actions should eventually evolve to zero if granted exhaustive training.) Finally, several units coded for the end-of-trial signal (Figure 4A,B) so that the network flushed the memories to prevent interference on subsequent trials.

The selectivity of example units has also been observed in animals. Figure 5A illustrates a cell in the lateral intraparietal area (LIP) in monkeys (Gottlieb and Goldberg, 1999) that transiently responded to one of the cues. Other neurons in monkey LIP persistently fire, thereby representing visual memory (Zhang & Barash, 2003) (Figure 5B). Similar activations can also be found in RECOLLECT units (see Figure 5C,D).

**Figure 5.**
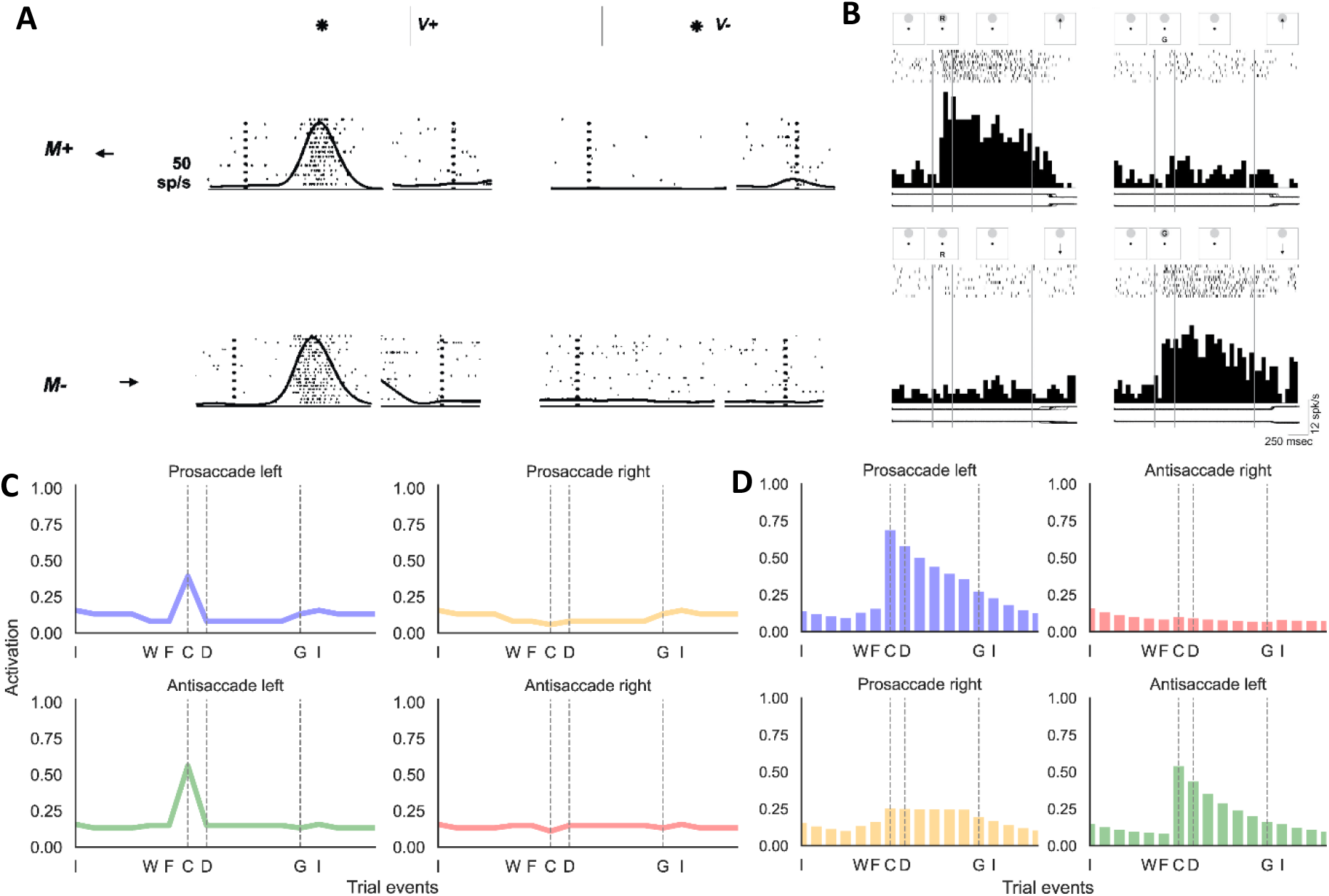
A) Example LIP neuron coding for a visual cue on the left (figure adapted from Gottlieb & Goldberg, 1999). B) Example LIP memory cell for cue location (Zhang & Barash, 2003). C) Candidate input unit in RECOLLECT coding for the left cue. D) Memory unit in RECOLLECT. Labels: I = intertrial interval, W = waiting period until fixation is acquired, F = time of fixation, C = cue presentation, D = memory delay period, G = go.

Other neurophysiological studies demonstrated that the duration of the persistent activity depends on the length of the memory period. When the memory delay is extended the memory activity of LIP neurons persists longer (Gnadt & Andersen, 1988) (Figure 6A).

**Figure 6.**
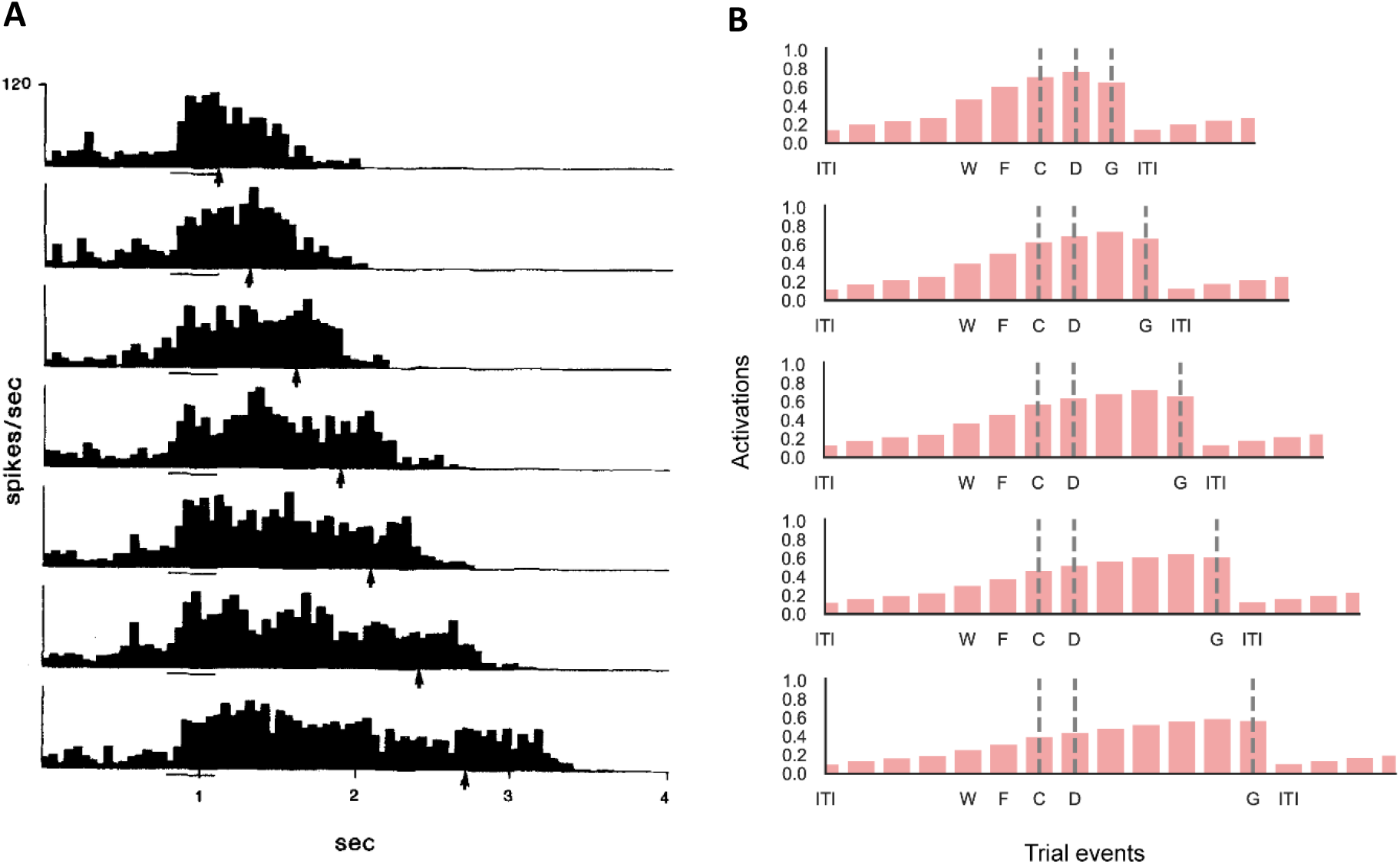
A) Neurons in lateral interparietal cortex (LIP) persistently fire for the length of the memory delay of the pro-/anti-saccade task (Gnadt & Andersen, 1988). B) Memory units in RECOLLECT also exhibit persistent firing across increasingly long delays (1, 2, 3, 4 or 5 timesteps, respectively). Labels: ITI = intertrial interval, W = waiting period until fixation is acquired, F = time of fixation, C = cue presentation, D = memory delay period, G = go.

To investigate whether RECOLLECT displays a similar behaviour, we trained a network with varying memory delays (one to five timesteps) (Figure 6B). The duration of persistent activity depended on the length of the delay, after which it declined; likely due to the end-of-trial signal.

We conclude that RECOLLECT can train networks on the pro/anti saccade-task. These networks learn to memorize and forget when necessary and use persistent activity to code for memories in a similar manner as neurons in the brain.

### RECOLLECT exhibits learning-to-learn on a reversal bandit task

We next investigated learning-to-learn with a reversal bandit task. This task has been used to assess meta-learning, including in Wang et al. (2018). Here, we use an architecture based on light-gated recurrent units trained using a biologically plausible learning rule – RECOLLECT – in order to elucidate the neural mechanisms underlying reversal learning (see Figure 7A). During the task, there are two levers that can be pulled. The two levers were randomly assigned a high (75%) or low (25%) reward probability. The task consisted of episodes with 100 discrete lever pulls, after which the reward probabilities were either reversed (reversal bandit), or randomly reassigned (random reversal bandit). The network had to sample the levers to assess which one yielded the higher reward and then harvest rewards by consistently pulling this lever until the end of an episode. The reversal bandit is easier than the random reversal bandit because the network can exploit the predictable reversal between successive episodes.

**Figure 7.**
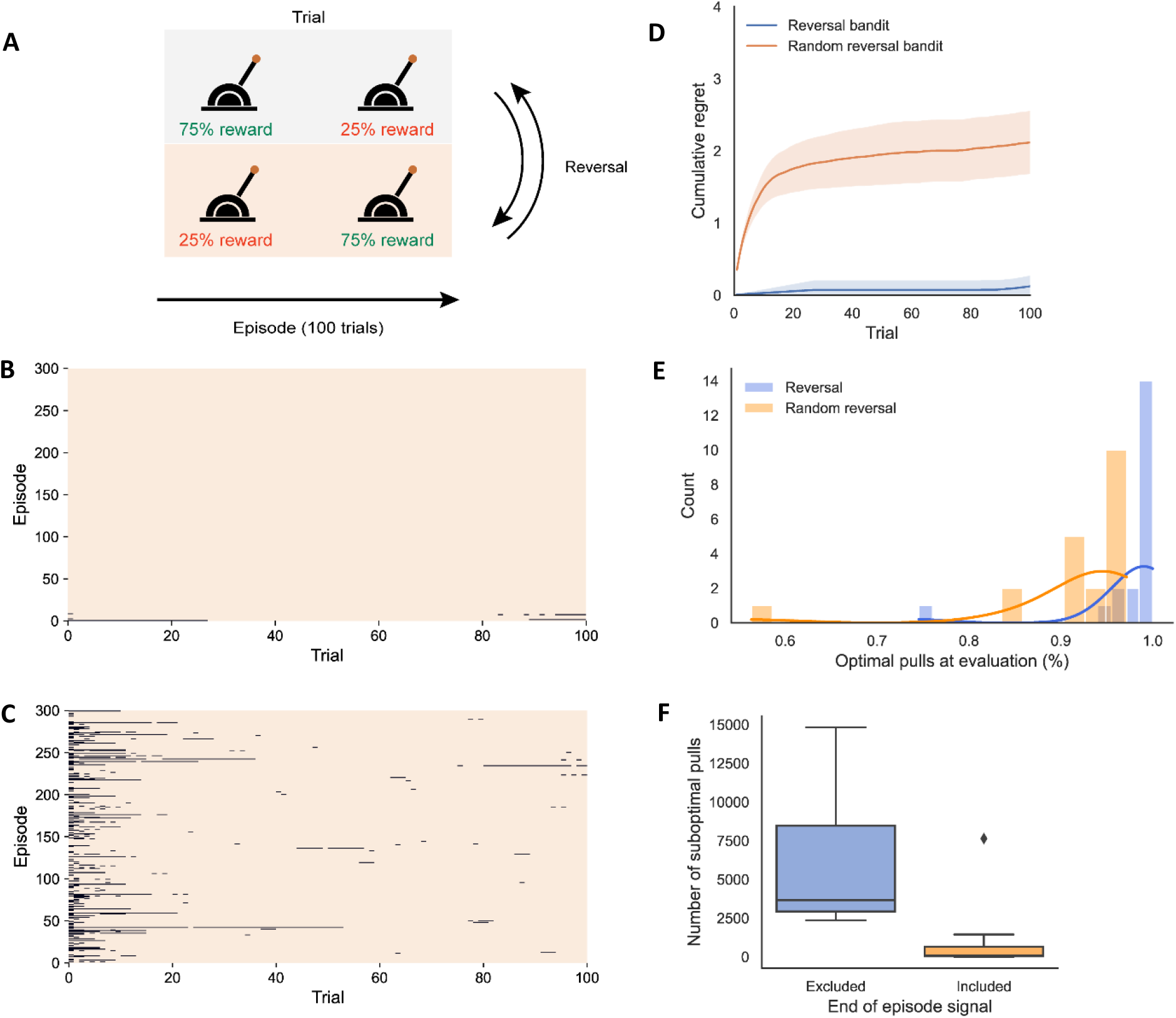
A) Structure of the two-armed bandit reversal task. In the case of the random version, a new episode is chosen randomly. B-C) Performance on an example network after training on the reversal bandit (B) and random reversal bandit (C) at evaluation (99.8% and 97.2% optimal pulls, respectively). The networks were initialised with the same seeds. Orange and black regions denote optimal and suboptimal choices, respectively. D) Cumulative regret (± 95% confidence interval) on (random) reversal bandit task. E) Histogram of the percentage optimal pulls on evaluation trials of the reversal bandit and random reversal bandit for the same 20 random seeds. F) Number of suboptimal pulls on the non-random reversal bandit is significantly lower when an end of episode signal is included.

Successful meta-learning on this task implies that a trained model can quickly (i.e. within one or just a few trials) switch to the new context at the start of a new episode by associating each context with a memory state, as opposed to relearning its weight structure to accommodate for the changed environment; something that would require hundreds of trials. To facilitate meta-learning, the network had access to the action it took on the previous timestep, and the reward it received. We also provided a signal that an episode had ended.

We trained RECOLLECT (with 4 input units, 4 gating, candidate input and memory units each (5 for the random reversal bandit), as well as 2 output units) on both versions of the reversal bandit task by presenting it with 20000 episodes of 100 trials each (as in Wang et al., 2018). Once the training phase was completed, learning and exploration were disabled and the model completed an additional 300 evaluation episodes. We evaluated performance as the number of choices of the low-rewarding (i.e. suboptimal) lever on 20 random initialisations of the network. For comparison with Wang et al. (2018), we also provide a measure of cumulative regret. Regret occurs when the action taken deviated from the optimal action (under hindsight) and a reward is not obtained. Cumulative regret refers to the cumulative loss of these expected rewards over time (Pepels et al., 2014).

Figures 7B,C illustrate the suboptimal pulls during the evaluation phase of two networks that were trained on the reversal and random reversal bandit tasks. The example network trained on the reversal bandit task learned to select the correct lever upon episode reversals almost perfectly. Suboptimal pulls only occurred either at the beginning of an episode or just before the end (perhaps to sample whether a new episode had already begun). On the random variant, there were more suboptimal arm pulls. These were concentrated at the beginning of episodes. While RECOLLECT tended to select the correct lever thereafter, it was not uncommon to observe exploratory actions being taken along the entire course of the episodes. Averaged over evaluation episodes, performance of RECOLLECT on the random reversal bandit (see Figure 7D) was comparable to – but slightly below – that of long-short term memory-based architectures, with an average cumulative regret of 2.1 for RECOLLECT (97.2% optimal pulls) versus 1.1 in Wang et al. (2018; 98.5% optimal pulls) on the same task with identical reward distributions, despite its reduced computational complexity and use of a local learning rule.

Networks trained on the reversal bandit task (see Figure 7E) achieved a median accuracy of 99.7%, with some networks reaching 100% optimal pulls. As expected, the accuracy on the random reversal bandit was lower with a median fraction of optimal pulls of 94.9%. A Wilcoxon Signed-Ranks test indicated that scores were significantly lower on the random reversal bandit compared to the non-random reversal bandit, *z* = -3.06, *p* = .002. This constituted a large effect, *r* = 0.68. These results indicate that RECOLLECT was able to exploit the additional regularity in the structure of the reversal bandit. Because the episodes alternated, the network learned to reverse its actions upon the end-of-episode signal and it did not have the sample the new reward structure when a new episode started.

To investigate the effect of the end-of-episode signal, we trained 20 networks with and without this signal on the non-random reversal bandit (Figure 7F). At evaluation, the median number of suboptimal pulls in the presence of the end-of-episode signal was 99, which was significantly lower than the 3661 pulls without this signal (Wilcoxon signed-ranks test, *z* = -3.47, *p* < .001), with a large effect size (*r* = .78). Hence, RECOLLECT capitalises on the end-of-episode signal to increase its performance.

We next analysed the activity of units in a network with the smallest size possible to perform the task (the hidden layer consists of two gating, candidate input and memory units each), to gain additional insight into how it solves the reversal bandit task. We plotted the average activation (± *SEM*) of all units in the network for left and right high-rewarding episodes before and after reversals for an example network (Figure 8). We will first discuss how these activations develop in the absence of an end of episode signal (see Figure 8A), after which the same analysis will be conducted with the signal included (Figure 8B).

**Figure 8.**
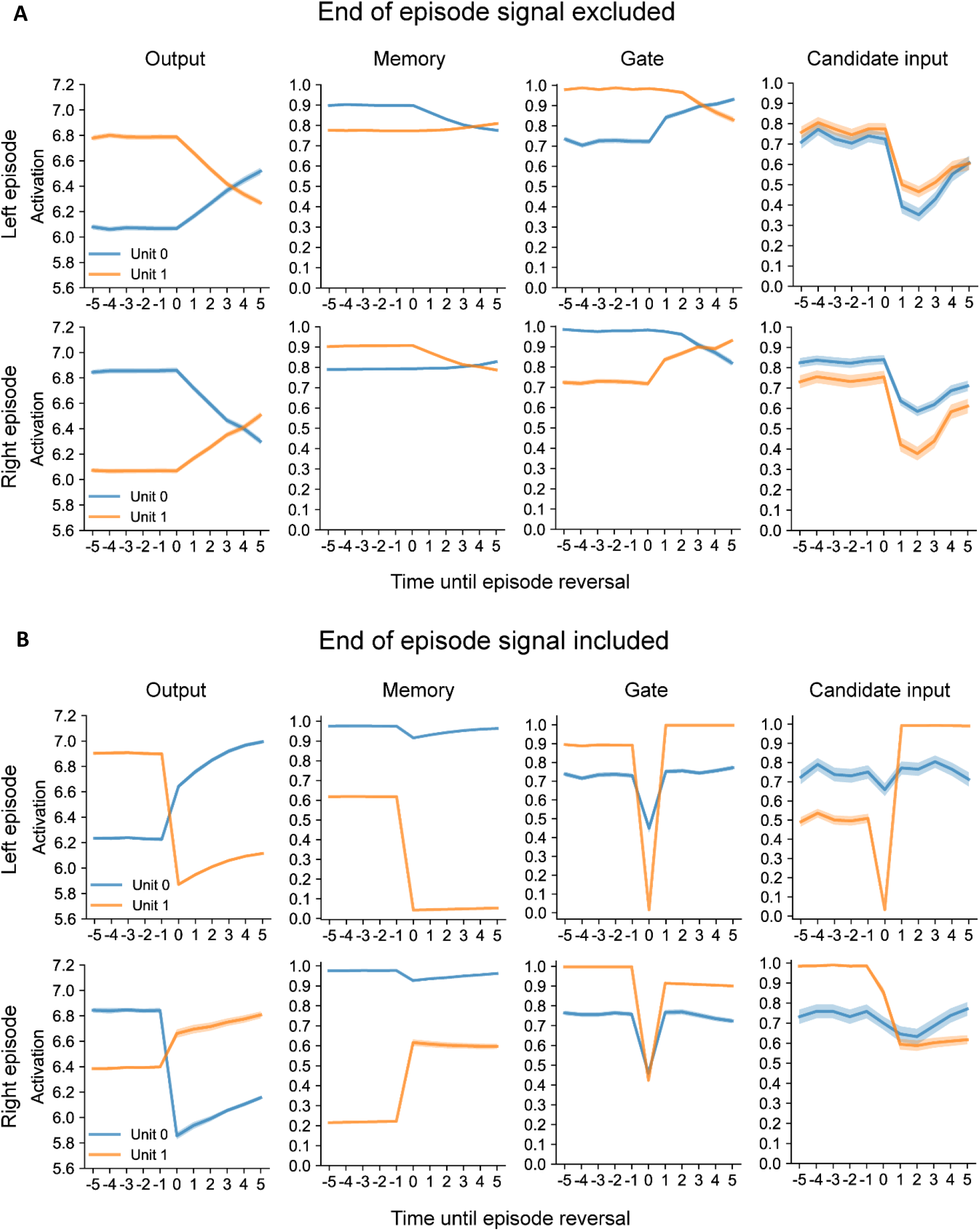
The average activity (± *SEM*) of all units in the network for left and right high-rewarding episodes on the non-random reversal bandit in the time window around the episode reversal. Activations are shown when no end of episode signal was provided (A) and when this signal was included (B). Note that the activities of the two output units reverse after the end of the episode. The reversal is faster if the end-of-episode signal is provided than if it is not. In the second column, it can also be seen how the memory units code of the task context. The end-of-trial signal gives rise to an abrupt change in the memory state, whereas the network has to gather evidence across trials based on the reward contingency that thecontext changed of theend-of-trial signal is absent. This is since only 75% of the optimal choiceare rewarded, one or two non-rewarded trials do not imply that the reversal has occurred. The third column shows that the end-of-trials signal causes a flushing of the memory state, which is visible as a reversal (A) or strong decrease in the activity of the gating unit (B). In the fourth column, candidate input units show reductions in activity upon episode switches and reverse their activations thereafter.

As expected, the activity of the Q-value unit coding for the highly rewarded action was higher than that of the other output unit. This pattern started to slowly reverse after the episode switch (*t* = 0) until the other unit was more highly activated (approximately at *t* = 4). This indicates the accumulation of evidence in these neurons about which context is relevant. Since the correct lever is only rewarded 75% of the time and the incorrect lever yields a reward 25% of the time, a single rewarded lever pull does not give reliable information about the context. Instead, RECOLLECT needs to integrate outcome information for a few trials until it is certain that the episode has changed.

A similar pattern can be observed for the memory units, where units coding for episode type reverse their activations along a comparable time course. For gating units this behaviour can also be seen. Interestingly, the activation of the gating unit selective for the current episode type is almost set to the maximum value of 1. Since high values for the gating units mean that all memory is maintained, this means that the gating units slowly allow for new content from the candidate input units to enter the memory upon the end of an episode; effectively initiating the reversal in activations for memory units.

While the two candidate units also show selectivity for episode type, these activations do not show a reversal following the same time course as other units. Instead, both units sharply decrease their activity (*t* = 1), perhaps due to the absence of the expected reward directly after the change in context, followed by a slower recovery.

Thus, due to a combination of the gating units slowly releasing content from the memory units that can then be supplemented by the episode-specific encodings from the candidate input units, the memory units – and by proxy the output units – can reverse their activations to accommodate for the episode changes in the absence of explicit reset signals.

In the case that an end of episode signal was provided to the network (see Figure 8B), the switch in output unit activity was accelerated, which indicates that the network learned the significance of this signal and uses it to efficiently (immediately at *t* = 0) change its activations to select the new optimal lever. However, the precise strategy how this is accomplished by the hidden units differs. Rather than having two memory units each selective for a different episode type and slowly reversing, one memory unit did not change its activity across episodes (unit 0, this unit was almost always fully activated, except for a slight decrease at the end of episodes) and the other unit showed sharp decreases and increases upon episode reversals for left and right episodes, respectively (unit 1).

For gating units, one unit (unit 1) was usually more active than the other across episode types. Moreover, it was even fully activated during right high-rewarding episodes. While both units showed steep declines in activation for both episode types upon reversals, unit 1 even almost completely flushed memory (activation is approximately 0), when the high-rewarding lever switched from left to right.

As with memory and gating units, one of the candidate input units (unit 0) did not exhibit much variation in activation over episode types. However, the other unit (unit 1) was almost fully active during right high-rewarding episodes, and had lower activity than the other unit during left high-rewarding episodes. Moreover, it showed a strong reduction in activation upon reversals (*t* = 0), particularly when the left lever became the low-rewarding lever. As a result, a reversal in activations occurred.

In summary, this indicates that rather than slowly accumulating evidence, RECOLLECT rapidly switches between memory states when an end of episode signal is provided. This is facilitated by strong flushing of memory by the gating units immediately upon episode reversal and concurrent changes in candidate input unit activations that can then be incorporated into memory. Moreover, while the overall behaviour of the output units remained similar, the neural representation of the task appeared more efficient for the other units, with only one gating, candidate input unit and memory unit showing substantial task-related selectivity. Therefore, RECOLLECT can learn to leverage the end of episode signal to improve both efficiency and performance on the reversal bandit.

Finally, we compared the behaviour of networks trained with RECOLLECT during the non-random reversal bandit to the choices made by rats trained on a similar task. Brunswik (1939) trained rats on serial-reversal task on a T-maze, with two arms that were baited with different rewards. On the first 24 trials, one arm was always rewarded and the other arm was never rewarded. Rewards were reversed for the subsequent 16 trials. This was followed by several reversal episodes of 8 trials each, until the rats completed a total of 8 episodes. During the first episode the performance gradually increased (Figure 9A). The first reversal caused a sharp increase in errors, which then declined, a pattern that repeated for every reversal afterwards. Interestingly, the rats required fewer trials to accommodate the later switches, indicating that the rats learned-to-learn this task.

**Figure 9.**
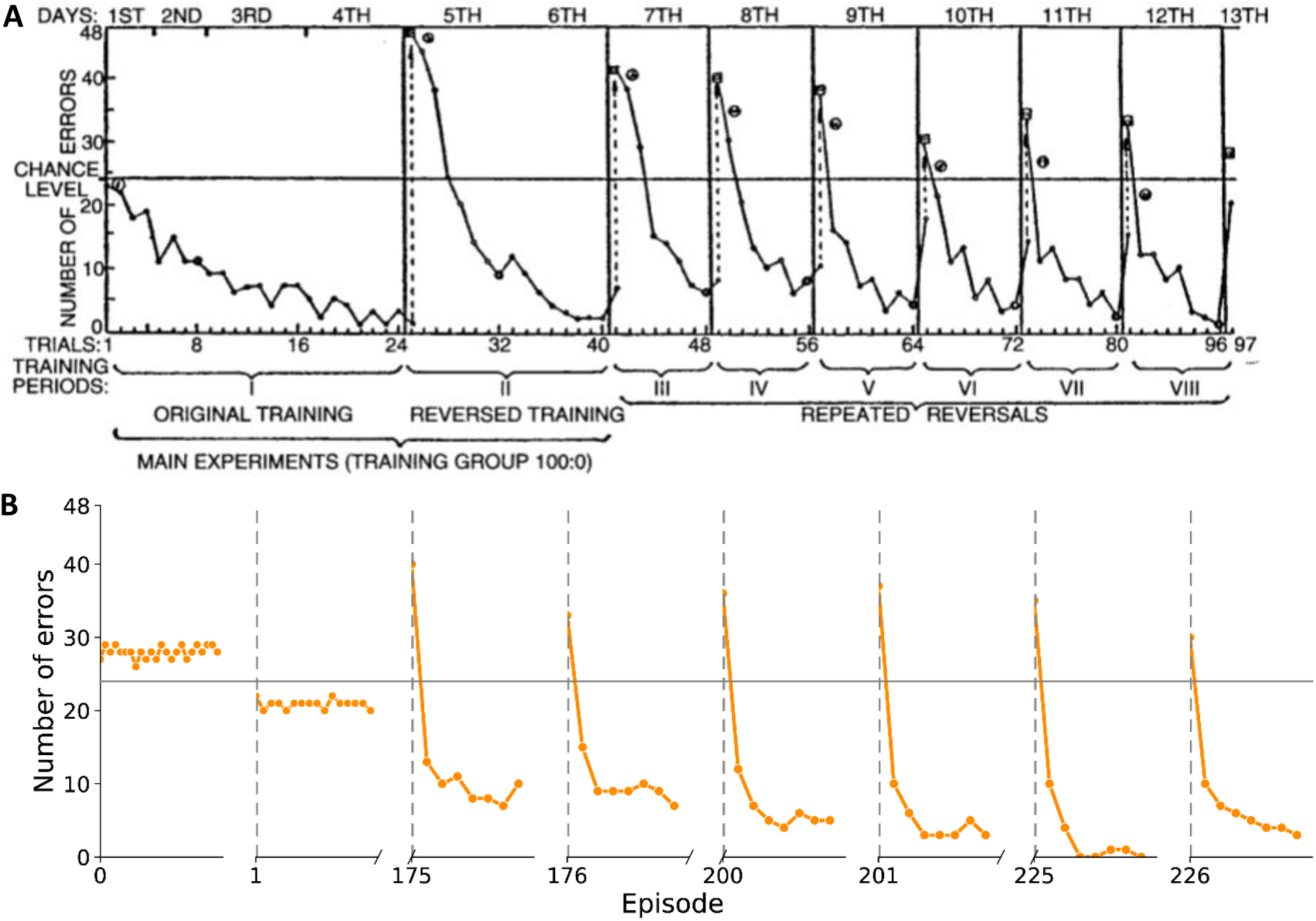
Comparison of errors made by networks trained with RECOLLECT and rats. (A) Learning on the reversal bandit task by rats (data from Brunswik, 1939) (B) Networks trained with RECOLLECT. We plotted the first 24 trials of the first episode, 16 trials after the first reversal and 8 trials of subsequent episodes.

We next analysed switching behaviour for 48 networks trained with RECOLLECT (Figure 9B) for the same reward distribution (i.e. 100% probability of receiving a reward for the high-rewarding lever and 0% probability of a reward for the low-rewarding lever). Since learning in RECOLLECT is slower than that of rats, we therefore plotted the number of errors in the first episode, the first reversal and the 175^th^, 200^th^ and 225^th^ episodes with the subsequent reversals (episode 176, 201 and 226). After this time, RECOLLECT showed a similar pattern as the rats: a relatively large number of errors upon the beginning of an episode, which decreased over the course of the episode. From the second episode onwards, the RECOLLECT data also showed the reduction in number of errors at the end of the episode over time similar to what was observed in Brunswik et al. (1939).

In conclusion, RECOLLECT can successfully train networks on the reversal bandit task in a way that is comparable to non-biologically plausible models. Moreover, the progression of learning is qualitatively similar to the behaviour of rats in a reversal task.

## Discussion

We developed a novel gated recurrent network that learned-to-learn in a biologically plausible manner.

The model incorporated the light-gated recurrent unit (Light-GRU; Ravanelli et al., 2018) and its learning rule was based on AuGMEnT (Rombouts et al., 2015). AuGMEnT is a learning method that uses a combination of attentional feedback and neuromodulators that code for the RPE to enable a form of learning that is similar to backpropagation-through-time, yet biologically plausible. In RECOLLECT, as in AuGMEnT, all information used to update the network is locally available at its synapses. Specifically, units in RECOLLECT can be considered part of the same cortical column and the feedback error as a locally available signal in that column. This signal might for instance be propagated by vasoactive intestinal peptide-positive (VIP+) interneurons, which are known to activate due to feedback and to gate learning (Zhang et al., 2014; Williams & Holtmaat, 2019). Hence, RECOLLECT uses a biologically plausible learning rule for gated recurrent units.

The main advantage of using a gated architecture is their flexibility. Whereas AuGMEnT remembers by default and can only forget by non-biologically plausible reset mechanisms that flush its entire memory (Rombouts et al., 2015), RECOLLECT can learn when and what to forget. Moreover, since the Light-GRU is a simplified version of more complicated gated architectures (Ravanelli et al., 2018), it is more easily interpretable and can therefore be used to elucidate neural mechanisms underlying memory and forgetting.

We tested RECOLLECT on a pro-/anti-saccade task, and found that the model flexibly selects which information to remember during a delay. Moreover, RECOLLECT learned to flush its memory at the end of a trial to prevent interference with subsequent trials. A comparison of units in networks trained with RECOLLECT to neurophysiological data revealed many similarities. Units developed selectivity for the colour of the fixation marker and the position of the cue, as well as persistent firing coding the relevant features, just as been observed in the visual and parietal cortex of monkeys (Gottlieb & Goldberg, 1999; Zhang & Barash, 2003; Gnadt & Andersen, 1988). Thus, RECOLLECT is not only biologically plausible given its reliance on neuromodulators and attentional feedback signals, but its units start to resemble neurons in the brains of animals that have learned the same task.

We used a reversal bandit task to test whether RECOLLECT learned-to-learn. Networks trained with RECOLLECT sampled the environment to gauge which of the two levers yielded the highest reward, and it then consistently chose this lever until the end of the episode. Moreover, the model’s behaviour during learning was reminiscent of how rats learn the reversal bandit task (Brunswik et al., 1939). There was an initial increase in errors upon the start of a new episode that decreased over the course of the episode. These errors declined more quickly as training progressed, indicating a similar progression of learning-to-learn in the model and in rats.

An interesting observation pertained to the role of the end-of-trial or end-of-episode signal. These signals enhanced performance by providing a signal that it is time to reconfigure the memory state to prepare for the upcoming episode. Likewise, cells in the frontal eye fields (FEF) have been shown to represent action sequence boundaries by increased firing rates following the end of the sequence (Fuji & Graybiel, 2003). Additionally, they supplied information about when the sequence would end, the peak response shifted to this moment in time, rather than shortly after the end of the sequence (Fuji & Graybiel, 2003). Similarly, Figure 8 depicts how the reversal in activations upon the end of an episode on the reversal bandit occurs faster in the presence of such a signal compared to when an end of episode signal was excluded. Therefore, RECOLLECT even parallels aspects of animal learning such as the identification of sequence boundaries.

RECOLLECT provides a model of how the brain can learn to memorize what is relevant and forget what is not, as was illustrated with the two tasks that we simulated. We note, however, that the precise mapping of the proposed mechanisms onto the circuits underlying memory and forgetting in the brain remains to be elucidated. Previous neuroscientific studies revealed multiregional loops between the cortex, thalamus and striatum for working memory (Bolkan et al., 2017; Wang et al., 2021; Schmitt et al., 2017; Rusu & Pennartz, 2020). Recent evidence also points towards a role of the loop through the cerebellum in working memory (de Zeeuw et al., 2021; Gao et al., 2018; Brissden et al., 2021; Schmahmann, 2019). These loops have also been implied in reversal learning (Parker et al., 2022; Tuite, Girotti, & Morilak, 2022). More research is needed to fully comprehend how these circuits effectuate working memory and forgetting.

One limitation of RECOLLECT pertains to its current architecture that consists of one layer with memory units. Future work could expand RECOLLECT so that it can be used in more complex tasks. Furthermore, while RECOLLECT consistently converged on the pro-/anti-saccade task, learning was slower than with other architectures such as AuGMEnT (Rombouts et al., 2016), which remember by default. Similarly, RECOLLECT performed less well on the random reversal bandit than LSTM-based networks. These differences are partially explained by the extra information that was given to the previous models. For example, in the study on AuGMEnT and in Wang et al. (2017), the network state was reset at the end of each trial. In a variant of AuGMEnT that had to learn to reset its working memory itself, learning was slower than in standard AuGMEnT (Rombouts et al., 2014). RECOLLECT learned the time structure of the task, what to remember and when to forget it. The network learned to take advantage of end-of-trial signals, but learning was even possible when such a signal was not presented.

There are also other learning rules and models that approximate backpropagation-through-time (e.g. Nicola & Clopath, 2017; Bellec et al., 2020). RECOLLECT uses the same approximation as e-prop (Bellec et al., 2020), which has been used to train long-short term memory models in reinforcement learning settings, and extends it to light-gated recurrent units. There are two main differences between RECOLLECT and e-prop. Firstly, RECOLLECT incorporates synaptic tags that implement the faster TD(λ) algorithm, rather than the simpler TD(0) method (van Seijen & Sutton, 2014). Secondly, e-prop requires each unit to be connected to an output unit to propagate the error signal. Since this is not biologically plausible for deeper networks, this property limits it to shallow networks, while RECOLLECT could be extended to more complicated architectures and tasks.

In conclusion, RECOLLECT is a novel gated recurrent neural network that by only using information available at the synapse can learn to flexibly use its working memory, as well as learn-to-learn in a manner that is reminiscent to that of animals. Therefore, it presents a biologically plausible alternative to traditional gated recurrent networks such as long-short term memory. Models like RECOLLECT can help to create insight into how working memory, forgetting and learning-to-learn are implemented by the brain.

## Materials & Methods

### Architecture details

#### Activation function

We used a sigmoid activation function for the gating units and candidate input units:

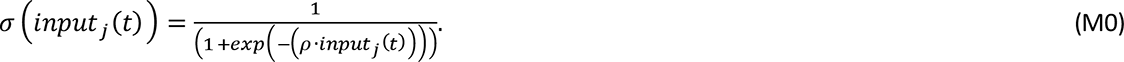

where *ρ* represents the slope of the sigmoid. This value was set to 2 in all experiments.

#### Learning rate

We noticed that rapid plasticity of gating units decreased the stability of learning. We therefore set the learning rate of synapse onto gating units was set at a lower value than those of other connections (see Table 1).

**Table 1.** RECOLLECT hyperparameters for each task (variant).

|  | Pro-/anti-saccade task | Reversal bandit | Random reversal bandit |
| --- | --- | --- | --- |
| Exploration rate ($\epsilon$) | 0.025 | 0.025 | 0.025 |
| Number of input units<br>(including end of<br>trial/episode signal) | 5 | 4 | 4 |
| Number of Light-GRU<br>units | 7 | 4 | 5 |
| Number of output units | 3 | 2 | 2 |
| Learning rate ($\beta$) | 0.1 | 0.01 | 0.005 |
| Learning rate of gating<br>units ($\beta_{gate}$) | 0.006 | 0.006 | 0.0005 |
| Discount factor ($\gamma$) | 0.9 | 0.9 | 0.9 |
| Tag decay rate ($\lambda$) | 0.4 | 0.2 | 0.1 |

#### Network parameters

A grid search with a limited set of a priori chosen values was conducted for parameter optimisation. Unless otherwise indicated, the parameters used for the experiments were as follows:

#### Pro-/anti-saccade task

To facilitate comparison with Rombouts et al. (2015), simulations regarding performance (Figure 3) on the pro-/anti-saccade task were performed using an intertrial interval of 1 and a memory delay of 2. In all further stimulations (except for Figure 6) we used an intertrial interval of 3 and memory delay of 5 timesteps so that the neural activations during memory delay and after the end of trial signal could be studied more closely. A curriculum was used to achieve these longer memory delays. Specifically, it involved using 1/5^th^ of the final memory delay in the beginning of the task. After the model reached criterion performance (85% correct trials on the previous 100 trials of each trial type), the current memory delay was multiplied by 2 until the final memory delay was reached (if this value exceeded this, it was set to the final memory delay). Networks contained 7 Light-GRU units (i.e. a set of one gating, candidate input and memory unit), except for Figure 5C,D, which were trained with a network size of 12 Light-GRU units.

#### Reversal bandit

In order to reach a better understanding of how RECOLLECT solves the reversal bandit, the activation plots and neural data comparison figure were created with networks containing the minimum network size possible to solve the task (i.e. two units each for the gating, candidate input and memory cells). For the performance figures please refer to the network size in Table 1.

#### Statistical analyses

Prior to statistical analysis, assumptions of normality were tested using the Kolmogorov-Smirnov and Shapiro-Wilk tests. If these tests indicated significant deviations from normality for at least one of the two distributions, a non-parametric test variant was used and the median was reported instead of the mean.

Cohen’s (1998) criteria were used to interpret effect sizes, where *r* = .1, *r* = .3 and *r* = .5 are considered small, medium and large effects, respectively.

### The relation between backpropagation-through-time and RECOLLECT

In this section, it will be demonstrated that backpropagation-through-time can be locally approximated in RECOLLECT using a combination of synaptic traces and tags.

#### Computing the gradient of M_j_(t), k_j_(t) and C_j_(t)

It follows from Fig. 1 that the influence of the activity of memory unit *j*, *M_j_(t)*, on the predicted value of the selected action *s*, *q_s_(t)*, is:

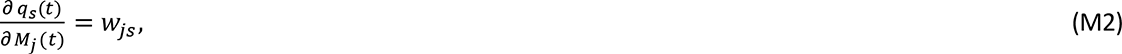

which is proportional to the amount of attentional feedback flowing from the winning action *s* to memory unit *j* (Roelfsema & van Ooyen, 2005). We can now compute the influence of the memory gate *k_j_(t)* on *q_s_(t)* using equation *M1*:

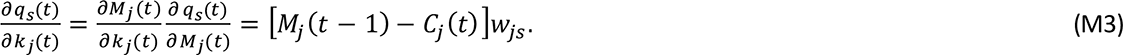

The influence of *C_j_(t)* on *q_s_(t)* depends on *k_j_(t)* and is as follows:

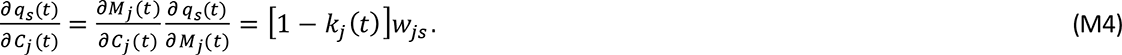

#### Computing the gradient of w^c^_ij_ using synaptic traces

We can now compute the *instantaneous* impact of connections *w^c^_*ij*_ (t)* on *q (t)*:

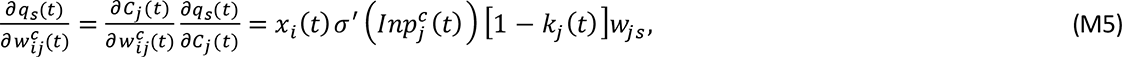

where 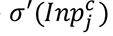 is the derivative of the activation function. However, these connections have also had impact on the memory state *M_j_(t)* on all previous time steps according to equation (1). For example, connection *w^c^_ij_*had an influence on *C_j_(t-1)* which influenced *M_j_(t-1)* and thereby also *M_j_(t)*. Although the notation is a bit ugly, for convenience let us write for this influence of *w^c^_ij_* on *t-1* on *q_s_(t)*:

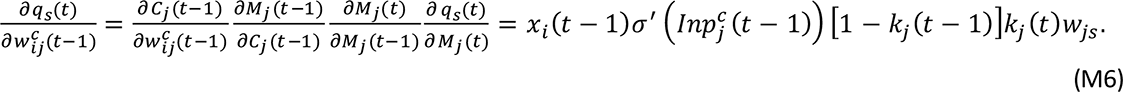

We can also compute this term for *t-2*:

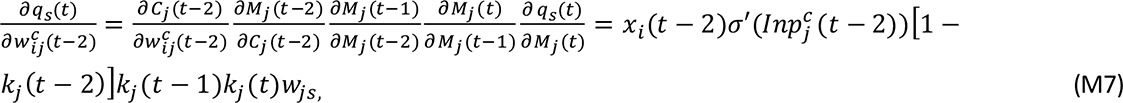

and, in general, for *t-i*:

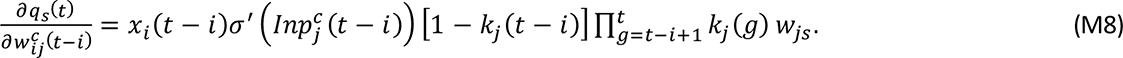

Although this gradient may look complex, it is actually straightforward to store the information in a *trace^c^_ij_* at the synapse and update it based on information that is locally available:

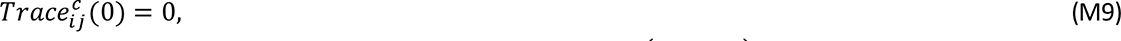

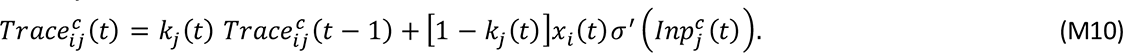

Adding all the time steps, the total influence of *w^c^_ij_* on *q_s_(t)* becomes:

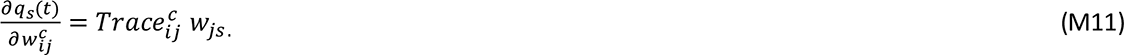

#### Computing the gradient of w^k^_ij_ using synaptic traces

Now it is time to use equation (5) to also compute the influence of the synapses *w^k^_ij_* that influence the memory gate *k_j_(t)* on *q_s_(t)*. As before, we start with the *instantaneous* impact of connections *w^k^_ij_(t)* on *q (t)*:

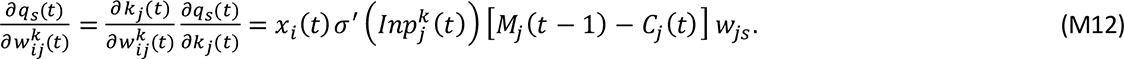

Let us now consider the influence of this synapse at *t-1* on the *q_s_(t)*:

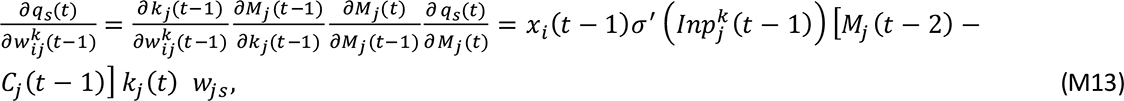

and at *t-2*

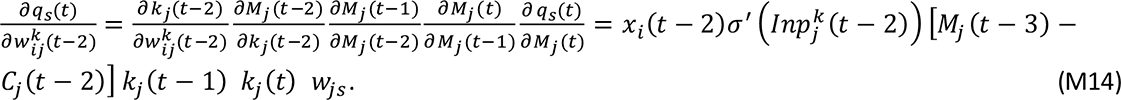

In general, for *t-i*:

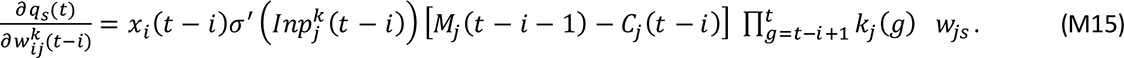

This gradient can also be stored in the form of a *trace^k^_ij_* at the synapse and updated based on information that is locally available:

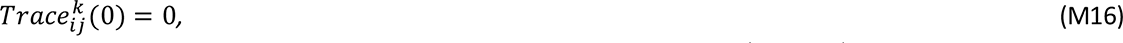

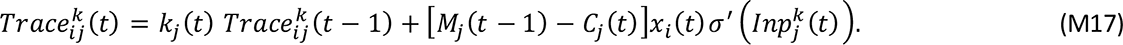

When adding all the time steps, the total influence of *w^k^_ij_* on *q_s_(t)* becomes:

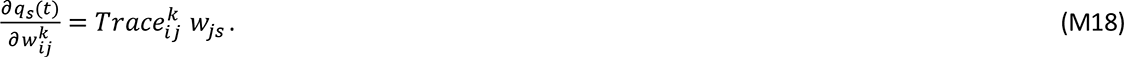

#### Tags and traces

RECOLLECT distinguishes between traces and tags (see also Rombouts et al., 2016). Whereas the traces represent the contribution of a synapse to the activity of the memory unit, the tags represent the influence of the synapse on the Q-value of the chosen action. The tag depends on the trace as well as on the amount of attentional feedback that arrives at the memory unit through the feedback connection from the chosen action (eq. 11 and 15).

The tags are used to implement the SARSA(λ) algorithm. If λ is larger than zero, the synapses that contributed to previous actions are also updated, while taking the temporal discount factor γ into account. This is an advantage of RECOLLECT over e.g. e-prop (Bellec et al., 2020), which uses the same approach to approximating backpropagation-through-time. The resulting combination of tags and traces (see equations 1-3), can be shown to be equivalent to gradient descent through backpropagation-through-time on the temporal difference error in the absence of recurrent connections (see Rombouts et al., 2015 for more detail), and to approximate backpropagation-through-time when recurrent weights are included, as in e-prop (Bellec et al., 2020).

## Acknowledgements

This research has received funding from the European Union’s Horizon 2020 Framework Programme for Research and Innovation under the Specific Grant Agreement No. 945539 (Human Brain Project SGA3, Task 3.7).

We acknowledge the use of Fenix Infrastructure resources, which are partially funded from the European Union’s Horizon 2020 research and innovation programme through the ICEI project under the grant agreement No. 800858.

## Footnotes

1 AuGMEnT was trained with 3 regular hidden units, 4 memory hidden units, 4 instantaneous input units and 8 transient input units (Rombouts et al., 2015). The total number of trainable weights was 75 for AuGMEnT and 94 for RECOLLECT.

## Notes

### Competing Interest Statement

The authors have declared no competing interest.

## References

Barraclough, D. J., Conroy, M. L., & Lee, D. (2004). Prefrontal cortex and decision making in a mixed-strategy game. Nature Neuroscience, 7(4), Article 4. https://doi.org/10.1038/nn1209

Bellec, G., Scherr, F., Subramoney, A., Hajek, E., Salaj, D., Legenstein, R., & Maass, W. (2020). A solution to the learning dilemma for recurrent networks of spiking neurons. Nature Communications, 11(1), Article 1. https://doi.org/10.1038/s41467-020-17236-y

Bolkan, S. S., Stujenske, J. M., Parnaudeau, S., Spellman, T. J., Rauffenbart, C., Abbas, A. I., Harris, A. Z., Gordon, J. A., & Kellendonk, C. (2017). Thalamic projections sustain prefrontal activity during working memory maintenance. Nature Neuroscience, 20(7), Article 7. https://doi.org/10.1038/nn.4568

Brissenden, J. A., Tobyne, S. M., Halko, M. A., & Somers, D. C. (2021). Stimulus-Specific Visual Working Memory Representations in Human Cerebellar Lobule VIIb/VIIIa. Journal of Neuroscience, 41(5), 1033–1045.

Brunswik, E. (1939). Probability as a determiner of rat behavior. Journal of Experimental Psychology, 25(2), 175–197. https://doi.org/10.1037/h0061204

Brunswik, E. (2001). The Essential Brunswik: Beginnings, Explications, Applications. Oxford University Press.

Cai, X., & Padoa-Schioppa, C. (2014). Contributions of Orbitofrontal and Lateral Prefrontal Cortices to Economic Choice and the Good-to-Action Transformation. Neuron, 81(5), 1140–1151. https://doi.org/10.1016/j.neuron.2014.01.008

Cho, K., van Merrienboer, B., Bahdanau, D., & Bengio, Y. (2014). On the Properties of Neural Machine Translation: Encoder-Decoder Approaches. ArXiv:1409.1259 [Cs, Stat]. http://arxiv.org/abs/1409.1259

Cohen, J. (1988). Statistical Power Analysis for the Behavioral Sciences (2nd ed.). Routledge. https://doi.org/10.4324/9780203771587

De Zeeuw, C. I., Lisberger, S. G., & Raymond, J. L. (2021). Diversity and dynamism in the cerebellum. Nature Neuroscience, 24(2), Article 2. https://doi.org/10.1038/s41593-020-00754-9

Dey, R., & Salem, F. M. (2017). Gate-variants of Gated Recurrent Unit (GRU) neural networks. 2017 IEEE 60th International Midwest Symposium on Circuits and Systems (MWSCAS), 1597–1600. https://doi.org/10.1109/MWSCAS.2017.8053243

Duan, Y., Schulman, J., Chen, X., Bartlett, P. L., Sutskever, I., & Abbeel, P. (2016). RL$^2$: Fast Reinforcement Learning via Slow Reinforcement Learning. ArXiv:1611.02779 [Cs, Stat]. http://arxiv.org/abs/1611.02779

French, R. M. (1999). Catastrophic forgetting in connectionist networks. Trends in Cognitive Sciences, 3(4), 128–135. https://doi.org/10.1016/S1364-6613(99)01294-2

Fujii, N., & Graybiel, A. M. (2003). Representation of action sequence boundaries by macaque prefrontal cortical neurons. Science (New York, N.Y.), 301(5637), 1246–1249. https://doi.org/10.1126/science.1086872

Gao, Z., Davis, C., Thomas, A. M., Economo, M. N., Abrego, A. M., Svoboda, K., De Zeeuw, C. I., & Li, N. (2018). A cortico-cerebellar loop for motor planning. Nature, 563(7729), Article 7729. https://doi.org/10.1038/s41586-018-0633-x

Gerstner, W., Lehmann, M., Liakoni, V., Corneil, D., & Brea, J. (2018). Eligibility Traces and Plasticity on Behavioral Time Scales: Experimental Support of NeoHebbian Three-Factor Learning Rules. Frontiers in Neural Circuits, 12, 53. https://doi.org/10.3389/fncir.2018.00053

Gnadt, J. W., & Andersen, R. A. (1988). Memory related motor planning activity in posterior parietal cortex of macaque. 70(1), 216–220. https://doi.org/10.1007/BF00271862

Gottlieb, J., & Goldberg, M. E. (1999). Activity of neurons in the lateral intraparietal area of the monkey during an antisaccade task. Nature Neuroscience, 2(10), 906–912. https://doi.org/10.1038/13209

Harlow, H. F. (1949). The formation of learning sets. Psychological Review, 56(1), 51–65. https://doi.org/10.1037/h0062474

Hochreiter, S., & Schmidhuber, J. (1997). Long Short-Term Memory. Neural Computation, 9(8), 1735– 1780. https://doi.org/10.1162/neco.1997.9.8.1735

Houk, J. C., Davis, J. L., & Beiser, D. G. (Eds.). (1995). A Model of How the Basal Ganglia Generate and Use Neural Signals That Predict Reinforcement. In Models of Information Processing in the Basal Ganglia. The MIT Press. https://doi.org/10.7551/mitpress/4708.003.0020

Huisman, M., van Rijn, J. N., & Plaat, A. (n.d.). A survey of deep meta-learning. Artificial Intelligence Review, 54(6), 4483–4541. https://doi.org/10.1007/s10462-021-10004-4

Ito, M., & Doya, K. (2009). Validation of Decision-Making Models and Analysis of Decision Variables in the Rat Basal Ganglia. Journal of Neuroscience, 29(31), 9861–9874. https://doi.org/10.1523/JNEUROSCI.6157-08.2009

Izquierdo, A., Brigman, J. L., Radke, A. K., Rudebeck, P. H., & Holmes, A. (2017). The neural basis of reversal learning: An updated perspective. Neuroscience, 345, 12–26. https://doi.org/10.1016/j.neuroscience.2016.03.021

Keller, G. B., & Mrsic-Flogel, T. D. (2018). Predictive Processing: A Canonical Cortical Computation. Neuron, 100(2), 424–435. https://doi.org/10.1016/j.neuron.2018.10.003

Khona, M., & Fiete, I. R. (2022). Attractor and integrator networks in the brain. Nature Reviews Neuroscience, 23(12), Article 12. https://doi.org/10.1038/s41583-022-00642-0

Kovach, C. K., Daw, N. D., Rudrauf, D., Tranel, D., O’Doherty, J. P., & Adolphs, R. (2012). Anterior Prefrontal Cortex Contributes to Action Selection through Tracking of Recent Reward Trends. Journal of Neuroscience, 32(25), 8434–8442. https://doi.org/10.1523/JNEUROSCI.5468-11.2012

Lillicrap, T. P., & Santoro, A. (2019). Backpropagation through time and the brain. Current Opinion in Neurobiology, 55, 82–89. https://doi.org/10.1016/j.conb.2019.01.011

Morris, G., Nevet, A., Arkadir, D., Vaadia, E., & Bergman, H. (2006). Midbrain dopamine neurons encode decisions for future action. Nature Neuroscience, 9(8), Article 8. https://doi.org/10.1038/nn1743

Nicola, W., & Clopath, C. (2017). Supervised learning in spiking neural networks with FORCE training. Nature Communications, 8(1), Article 1. https://doi.org/10.1038/s41467-017-01827-3

Padoa-Schioppa, C., & Assad, J. A. (2006). Neurons in the orbitofrontal cortex encode economic value. Nature, 441(7090), Article 7090. https://doi.org/10.1038/nature04676

Panichello, M. F., DePasquale, B., Pillow, J. W., & Buschman, T. J. (2019). Error-correcting dynamics in visual working memory. Nature Communications, 10(1), Article 1. https://doi.org/10.1038/s41467-019-11298-3

Parker, N. F., Baidya, A., Cox, J., Haetzel, L. M., Zhukovskaya, A., Murugan, M., Engelhard, B., Goldman, M. S., & Witten, I. B. (2022). Choice-selective sequences dominate in cortical relative to thalamic inputs to NAc to support reinforcement learning. Cell Reports, 39(7), 110756. https://doi.org/10.1016/j.celrep.2022.110756

Pepels, T., Cazenave, T., Winands, M. H. M., & Lanctot, M. (2014). Minimizing Simple and Cumulative Regret in Monte-Carlo Tree Search. In T. Cazenave, M. H. M. Winands, & Y. Björnsson (Eds.), Computer Games: Third Workshop on Computer Games, CGW 2014, Held in Conjunction with the 21st European Conference on Artificial Intelligence, ECAI 2014, Prague, Czech Republic, August 18, 2014, Revised Selected Papers 3 (pp. 1–15). Springer International Publishing. https://doi.org/10.1007/978-3-319-14923-3_1

Ravanelli, M., Brakel, P., Omologo, M., & Bengio, Y. (2018). Light Gated Recurrent Units for Speech Recognition. IEEE Transactions on Emerging Topics in Computational Intelligence, 2(2), 92–102. https://doi.org/10.1109/TETCI.2017.2762739

Roelfsema, P. R., & van Ooyen, A. (2005). Attention-gated reinforcement learning of internal representations for classification. Neural Computation, 17(10), 2176–2214.

Rombouts, J. O., Bohte, S. M., & Roelfsema, P. R. (2015). How Attention Can Create Synaptic Tags for the Learning of Working Memories in Sequential Tasks. PLOS Computational Biology, 11(3), e1004060.

Rombouts, J. O., Roelfsema, P. R., & Bohte, S. M. (2014). Learning Resets of Neural Working Memory. ESANN, 6.

Rushworth, M. F. S., Noonan, M. P., Boorman, E. D., Walton, M. E., & Behrens, T. E. (2011). Frontal Cortex and Reward-Guided Learning and Decision-Making. Neuron, 70(6), 1054–1069. https://doi.org/10.1016/j.neuron.2011.05.014

Rusu, S. I., & Pennartz, C. M. A. (2020). Learning, memory and consolidation mechanisms for behavioral control in hierarchically organized cortico-basal ganglia systems. Hippocampus, 30(1), 73–98. https://doi.org/10.1002/hipo.23167

Schmahmann, J. D. (2019). The cerebellum and cognition. Neuroscience Letters, 688, 62–75. https://doi.org/10.1016/j.neulet.2018.07.005

Schmitt, L. I., Wimmer, R. D., Nakajima, M., Happ, M., Mofakham, S., & Halassa, M. M. (2017). Thalamic amplification of cortical connectivity sustains attentional control. Nature, 545(7653), Article 7653. https://doi.org/10.1038/nature22073

Schultz, W. (2016). Dopamine reward prediction-error signalling: A two-component response. Nature Reviews Neuroscience, 17(3), 183–195. https://doi.org/10.1038/nrn.2015.26

Seijen, H., & Sutton, R. (2014). True Online TD(lambda). Proceedings of the 31st International Conference on Machine Learning, 692–700. https://proceedings.mlr.press/v32/seijen14.html

Sutton, R. S. (1988). Learning to predict by the methods of temporal differences. Machine Learning, 3, 9–44. https://doi.org/10.1007/BF00115009

Sutton, R. S. (2022). A History of Meta-gradient: Gradient Methods for Meta-learning. ArXiv:2202.09701 [Cs]. http://arxiv.org/abs/2202.09701

Sutton, R. S., & Barto, A. G. (2018). Reinforcement Learning, second edition: An Introduction. MIT Press.

Thrun, S., & Pratt, L. (1998). Learning to Learn: Introduction and Overview. In S. Thrun & L. Pratt (Eds.), Learning to Learn (pp. 3–17). Springer US. https://doi.org/10.1007/978-1-4615-5529-2_1

Wang, J. X. (2021). Meta-learning in natural and artificial intelligence. Current Opinion in Behavioral Sciences, 38, 90–95. https://doi.org/10.1016/j.cobeha.2021.01.002

Wang, J. X., Kurth-Nelson, Z., Kumaran, D., Tirumala, D., Soyer, H., Leibo, J. Z., Hassabis, D., & Botvinick, M. (2018). Prefrontal cortex as a meta-reinforcement learning system. Nature Neuroscience, 21(6), Article 6. https://doi.org/10.1038/s41593-018-0147-8

Wang, J. X., Kurth-Nelson, Z., Tirumala, D., Soyer, H., Leibo, J. Z., Munos, R., Blundell, C., Kumaran, D., & Botvinick, M. (2016). Learning to reinforcement learn. ArXiv:1611.05763 [Cs, Stat]. https://arxiv.org/abs/1611.05763v1

Wang, Y., Yin, X., Zhang, Z., Li, J., Zhao, W., & Guo, Z. V. (2021). A cortico-basal ganglia-thalamo-cortical channel underlying short-term memory. Neuron, 109(21), 3486–3499. https://doi.org/10.1016/j.neuron.2021.08.002

Williams, L. E., & Holtmaat, A. (2019). Higher-Order Thalamocortical Inputs Gate Synaptic Long-Term Potentiation via Disinhibition. Neuron, 101(1), 91–102. https://doi.org/10.1016/j.neuron.2018.10.049

Yamaguchi, K., Maeda, Y., Sawada, T., Iino, Y., Tajiri, M., Nakazato, R., Ishii, S., Kasai, H., & Yagishita, S. (2022). A behavioural correlate of the synaptic eligibility trace in the nucleus accumbens. Scientific Reports, 12(1), Article 1. https://doi.org/10.1038/s41598-022-05637-6

Zhang, M., & Barash, S. (2004). Persistent LIP Activity in Memory Antisaccades: Working Memory For a Sensorimotor Transformation. Journal of Neurophysiology, 91(3), 1424–1441.

Zhang, S., Xu, M., Kamigaki, T., Do, J., Chang, W.-C., Jenvay, S., Miyamichi, K., Luo, L., & Dan, Y. (n.d.). Long-range and local circuits for top-down modulation of visual cortex processing. Science, 345(6197), 660–665. https://doi.org/10.1126/science.1254126

